# Chemogenetic and optogenetic strategies for spatiotemporal control of split-enzyme-based calcium recording

**DOI:** 10.1101/2025.07.22.665990

**Authors:** Yunlei Zhang, Brianna K. Campmier, Akaash Sharma, Kelly N. Eckartt, Scott T. Laughlin

**Author notes:** Corresponding Author Correspondence should be addressed to Scott Laughlin.

## Abstract

Methods for monitoring physiological changes in cellular Ca^2+^ levels have been in high demand for their utility in monitoring neuronal signaling. Recently, we introduced SCANR (Split-Tobacco Etch Virus (TEV) protease Calcium-regulated Neuron Recorder), which reports on Ca^2+^ changes in cells through the binding of calmodulin and M13 to reconstitute an active TEV protease. First-generation SCANR marked all of the Ca^2+^ spikes that occur throughout the lifetime of the cell, but it did not have a mechanism for controlling the time window in which recording of physiological changes in Ca^2+^ occurred. Here, we explore both chemical and light-based strategies for controlling the time and place in which Ca^2+^ recording occurs. We describe the adaptation of six popular chemo- and opto-genetics methods for controlling protein activity and subcellular localization to the SCANR system. We report two successful strategies, one that leverages the LOV-Jα optogenetics system for sterically controlling protein interactions and another that employs chemogenetic manipulation of subcellular protein distribution using the FKBP/FRB rapamycin binding pair.

---

Neurons propagate electrical signals in expansive neural networks in the brain and peripheral nervous system. These networks rely on biochemical signals like Ca^2+^ flux to modulate cellular activity. For example, when neurons fire an action potential, they experience a dramatic increase in intracellular Ca^2+^ that leads to the release of neurotransmitters. The neurotransmitters bind to post-synaptic receptors on the receiving neuron resulting in signal propagation through the neural network. Because of the common physiology of neural signaling among the myriad types of neurons in the brain, methods to monitor Ca^2+^ changes serve as powerful reporters of neural activity.

There are several methods to mark neural circuitry with genetically encoded proteins that yield a time resolved change in fluorescence in response to a Ca^2+^ concentration spike. Perhaps the most popular are the GCaMP family of sensors. These are fusion proteins comprised of a Ca^2+^ binding domain and a circularly permuted GFP^1^. When in the presence of transiently high Ca^2+^ concentrations, the conformational change of the binding domain alters the beta barrel of GFP, increasing its fluorescence quantum yield^2^. This method has been used to great effect in the analysis of dynamic Ca^2+^ changes in model organisms such as zebrafish, Drosophila, *C. elegans*, and rodents^3,4^. However, in order to visualize activity throughout the entire brain, the organism under scrutiny needs to be small, the imaging speeds must be fast compared to those possible with conventional confocal microscopes, and the behavior or other neural system being studied must not be perturbed by the constrains of concurrent imaging and behavioral analysis.

In order to overcome these challenges, several groups have developed innovative methods for permanently marking active populations of neurons. For example, Schrieter and coworkers developed a permanent Ca^2+^ recording strategy called Calcium Modulated Photoactivatable Ratiometric Integrator (CaMPARI)^5^. Essentially, CaMPARI is a fusion between a Ca^2+^ binding domain and a green-to-red photoconvertible protein that photoconverts only in the presence of both high Ca^2+^ and photo-converting light. This method is a powerful tool for imaging whole brains with fewer limitations on imaging speed, brain size, and the types of behavior than can be studied. However, it is not without drawbacks, including the need to re-stimulate the fluorescent protein continuously and the inability to use optogenetics tools or alternative fluorescent reporters as the readout.

In another strategy, Luo and coworkers produced a system for permanent recording of Ca^2+^ spikes based on a transcriptional reporter, TRIC^6^. This system employs a calmodulin-linked Gal4 DNA binding domain, which, in the presence of high Ca^2+^, recruits a transcription activation domain to a gene of interest leading to expression of a user-defined effector protein, such as GFP. Although TRIC has been successfully employed in Drosophila, it requires modifications to work within different cell types due to its reliance on nuclear Ca^2+^, which varies between cell types, complicating its use in full brain mapping or in other popular model systems. Further, TRIC activation requires prolonged high levels of Ca^2+^ and can be saturated at basal Ca^2+^ levels.

Recently, Ting and coworkers described a clever genetic system for marking active neuron populations^7^. Called FLARE, this system combines Ca^2+^-regulated protein recruitment and blue-light-modulated substrate exposure to TEV protease. Essentially, FLARE employs a membrane-bound calmodulin-binding peptide linked to an evolved LOV (eLOV) domain, a TEV protease cleavage site, and a transcriptional factor. Upon exposure to high Ca^2+^ and blue light, the calmodulin-bound TEV protease is recruited to the cell membrane and the TEV protease cleavage site is revealed by the light-induced conformational change of the eLOV domain. This dual control leads to high signal-to-noise and negligible dark state leak, and the utilization of a transcription factor enables the use of a wide range of reporters. However, this system required several genetic optimizations and modifications to move between cultured system HEK293 cells and rat neurons, has a significant (hours) time-scale of the final readout of the system (though the activation of the system may occur within minutes), and required sustained *in vitro* stimulation of neurons for 15 min to achieve the reported signal levels^7^.

Finally, we recently reported a permanent Ca^2+^ recording system called the Split TEV Calcium-regulated Active Neuron Recorder (SCANR). SCANR marks Ca^2+^ concentration spikes with a user-defined signaling output, such as translocation of a pro-fluorescent protein or the translocation of a transcription activator like Gal4 that can lead to the downstream expression of a protein of interest^8^ (**Figure 1A**). This system responds to Ca^2+^ concentration spikes by dimerizing two Ca^2+^ binding domains, calmodulin and M13, resulting in the reformation of the split N- and C-termini of TEV protease. Reconstituting the enzyme restores its activity so that it can act upon its substrate to produce a trackable signal.

**Figure 1.**
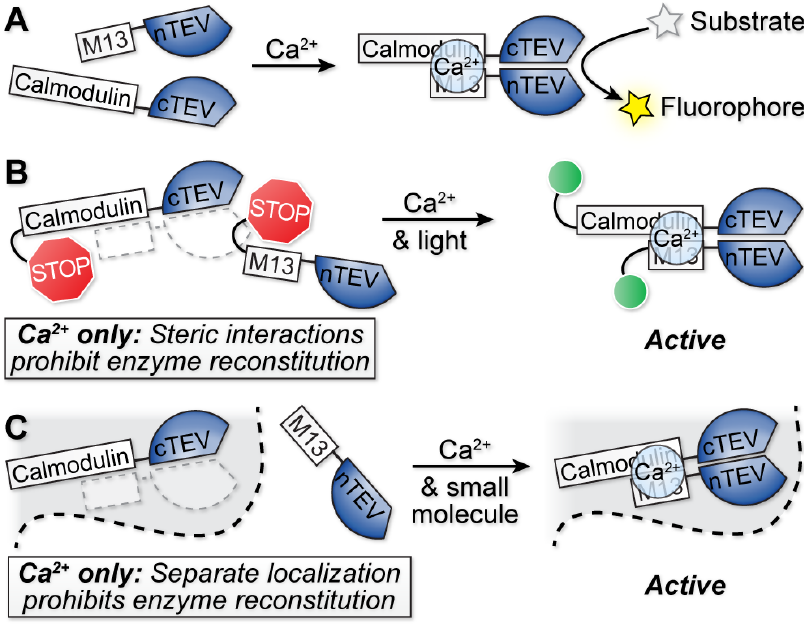
Strategies to control SCANR-based Ca^2+^ recording with light or small molecules. (**A**) Unmodified SCANR consists of two halves of a split Tobacco Etch Virus (TEV) protease linked to the Ca^2+^-binding domains calmodulin and M13. In this original version of SCANR, the split enzyme becomes active in the presence of high Ca^2+^ leading to the reconstitution of an active TEV protease that can cleave a genetically encoded substrate to mark Ca^2+^ concentration spikes in cells. (**B**) Steric blocking of the SCANR components using optogenetics switches would permit Ca^2+^ recording to occur only when desired by the experimenter. (**C**) Sequestering the SCANR pairs in different subcellular compartments prevents their association in the presence of high Ca^2+^. Inducing translocation of a SCANR component to a shared subcellular compartment by exposure to light or a small molecule provides control over the time window of Ca^2+^ recording.

The first iteration of SCANR measured all Ca^2+^ events in the lifetime of the cell after SCANR’s expression. That is, there was no built-in method to control the desired recording time window. Fortunately, the split protein nature of SCANR provides multiple avenues for user defined control of its function. Here, we describe optogenetic and chemogenetic approaches for controlling SCANR-based Ca^2+^ recording using on/off steric blocking of split protein reconstitution (**Figure 1B**) or manipulation of subcellular localization to control SCANR recording (**Figure 1C**). More specifically, we explore LOV-Jα-based control of both key epitope steric blocking and subcellular localization, CRY2-C1B1 optogenetic control of subcellular localization, and small molecule (rapamycin)-based control of subcellular localization.

## MATERIALS AND METHODS

### Cloning of modified SCANR and control expression vectors

The protocols and materials used to create the plasmids used in this study are detailed in the Supporting Information.

### Ionomycin-based assay for transiently increasing intracellular Ca^2+^ in HEK293T cells

These assays were performed essentially as described previously^8^. Briefly, cells were washed once with PBS and then incubated in 5 μM ionomycin in Hank’s balanced salt solution (HBSS) at room temperature for 5 min (when the total volume was 400 μL) or 8 min (when the total volume 2 mL). After the incubation, the cells were washed with phosphate buffered saline (PBS) to remove traces of ionomycin, placed into pre-warmed (37 °C) Dulbecco’s Modified Eagle Medium (DMEM) supplemented with 10% fetal calf serum, 100 U/ml penicillin, and 0.1 mg/ml streptomycin, and incubated overnight at 37 °C in air with 5% CO_2_.

### Blue light exposure protocols for optogenetics

Blue light stimulation for CRY2/C1B1was performed on live cells using a Zeiss Axio Vert.A1 inverted microscope equipped with a Zeiss EC Plan-Neofluor 20x NA0.50 objective. The green channel excitation was 480/40. Cells were illuminated with blue light for no more than 3 s per field of view. The illumination was performed both before and halfway through the ionomycin incubation procedure.

Blue light stimulation for the LOV constructs was performed on live cells using a Zeiss Axio Examiner.D1 modified with an Andor Differential Scanning Disk 2 confocal unit equipped with a 10x NA0.3 air objective and piezo objective holder for acquiring z-stacks. Optimal z-stack step size of 0.38 μm was calculated using the Andor iQ3 software to provide Nyquist sampling. The excitation filter used was 390/40 at 75% intensity at 200 ms for 12 repeats (30 s intervals) for 6 min.

Blue light stimulation for the LEXY, LINuS, and LANS constructs were performed on live cells using a Zeiss Axio Examiner.D1 modified with an Andor Differential Scanning Disk 2 confocal unit equipped with a 40x NA1.0 water immersion objective and piezo objective holder for acquiring Z-stacks.

Optimal z-stack step size of 0.38 μm was calculated using the Andor iQ3 software to provide Nyquist sampling. The excitation filter used was 390/40 at 100% intensity with a 40 ms exposure for 80 repeats (30 s intervals) for 40 min (LEXY), 1 s exposure for 30 repeats (30 s intervals) for 15 min (LINuS), and 300 ms exposure for 300 repeats (5 s intervals) for 25 min (LANS).

### Rapamycin assays for FKBP/FRB dimerization

The media was removed from cells expressing the rapamycin-inducible SCANR constructs and substrate and these cells were incubated with the indicated rapamycin concentration (100 nM–10 μM rapamycin) in DMEM for the indicated duration prior to exposure to ionomycin by addition of 5 μM ionomycin/rapamycin/HBSS. After the 5 min ionomycin exposure, the cells were washed with 400 μL PBS, pH 7.4, and incubated in the indicated [rapamycin] in DMEM supplemented with 10% fetal calf serum, 100 U/ml penicillin, and 0.1 mg/ml streptomycin, for 24 h at 37 °C in air with 5% CO_2_.

### Confocal imaging and analysis

Confocal microscopy was performed using a Zeiss Axio Examiner.D1 modified with an Andor Differential Scanning Disk 2 confocal unit equipped with a 40x NA1.0 water immersion objective and piezo objective holder for acquiring z-stacks. Optimal z-stack step size of 0.38 μm was calculated using the Andor iQ3 software to provide Nyquist sampling. Excitation and emission filters used were: Blue channel excitation 390/40, emission 452/45; Green channel excitation 482/18, emission 525/45; Red channel excitation 556/20, emission 609/54; Far red channel excitation 640/14. Emission 676/29. Confocal images were converted to maximum intensity projections of the z-stack and level-normalized across all images using ImageJ. Image cropping and organization into figure illustrations was performed using Adobe Illustrator CC. Fixed cells were randomly sampled for each condition. A maximum intensity z-projection of the red channel confocal image for each cell was created containing the slices with the nucleus. An ROI was drawn around the nucleus of each cell and the mean intensity of the pixels in that region was collected and exported to Microsoft Excel for graphing and statistical analyses (Two-tailed Student’s t-test).

## RESULTS AND DISCUSSION

The initial characterization of SCANR^8^ and previous reports of other split-TEV protease systems^9^ showed that expressing the two TEV protease halves in separate subcellular locales prohibited reformation of the active enzyme and turnover of the substrate. Based on these findings, one strategy for controlling SCANR Ca^2+^ recording is to segregate each part of the system into distinct subcellular compartments until induced to translocate to a shared location by an external stimulus. The other general strategy we explored takes advantage of optogenetic conformational switches that transition between protein conformations and are designed to either block or permit protein association between the individual SCANR components.

Exploring application to SCANR of LOV-Jα-based methods for optogenetic control of subcellular localization (LEXY, LINuS, and LANS). We began this study by exploring several methods for manipulating subcellular localization to control SCANR Ca^2+^ recording using the LOV-Jα optogenetic system. The LOV-Jα domain is an optogenetics workhorse that has been employed extensively to manipulate subcellular localization^10–12^ using clever methods like LEXY^10^, LINuS^13^, and LANS^11^. LEXY, LINuS, and LANS shuttle protein fusion cargo between subcellular compartments by taking advantage of nuclear import and export sequences that are exposed or hidden based on the conformational state of LOV-Jα. These tools differ in the details and directionality of transport: LEXY is a nuclear exporter, while LINuS and LANS are nuclear importers. Each method has been utilized to move a variety of proteins to the different subcellular locations,^10,11,13^ and the apparent ease with which different enzymes and proteins were plugged into these systems made them a tempting first choice for engineering optogenetic control into SCANR.

We explored the application of LEXY, LINuS, and LANS to controlling SCANR by fusing the C- or N-terminal component to the appropriate optogenetics cassette. We used the reported LEXY constructs^10^ and chose from the variety of available constructs based on the strength of the NLS and NES translocation sequences utilized for the reported LINuS and LANS variants^11,13^. We expressed these constructs in HEK293 cells, used the reported plasmids as controls to develop an appropriate illumination protocol for translocation, and assayed for translocation of the relevant SCANR fusions between the nucleus and cytoplasm.

Unfortunately, in each case, the subcellular distribution of the SCANR constructs was insufficient to permit control of their association and reactivity. For example, the SCANR constructs based on the LEXY system, which functions by hiding or exposing a nuclear export signal in response to blue light (**Figure 2A**), produced constructs with incomplete cytoplasm occlusion and modest nuclear to cytoplasm translocations. Importantly, the reported constructs showed a 2.2-fold decrease in nuclear levels of fluorescence after exposure to blue light (**Figure 2B**), similar to the 3-fold decrease reported in the literature for LEXY constructs with an appended flurophore^10^. However, several LEXY-modified SCANR constructs produced less impressive results. A LEXY-modified M13-SCANR construct (SCANR^M^-LEXY) had a roughly equal distribution between the cytoplasm and nucleus in the dark state, and a very modest 1.3-fold decrease in nuclear fluorescence levels after exposure to blue light (**Figure 2C**). A LEXY-modified calmodulin-SCANR construct (SCANRC*-LEXY), had a more respectable 1.8-fold change in nuclear fluorescence after exposure to blue light, but it also possessed noticeable cytoplasmic localization in the dark state, which would likely contribute to background Ca^2+^ recording (**Figure 2D**). Finally, two other LEXY-modified calmodulin SCANR constructs with either an mCherry (SCANR^C^-mCh-LEXY) or EGFP fluorophore (SCANR^C^-EGFP-LEXY) possessed very high cytoplasmic fluorescence in the dark state (**Figure 2E, F**). None of these constructs produced clear, low-level cytoplasmic localization in the dark state and high levels of cytoplasmic localization after exposure to blue light, making them unlikely to be useful for controlling SCANR activation.

**Figure 2.**
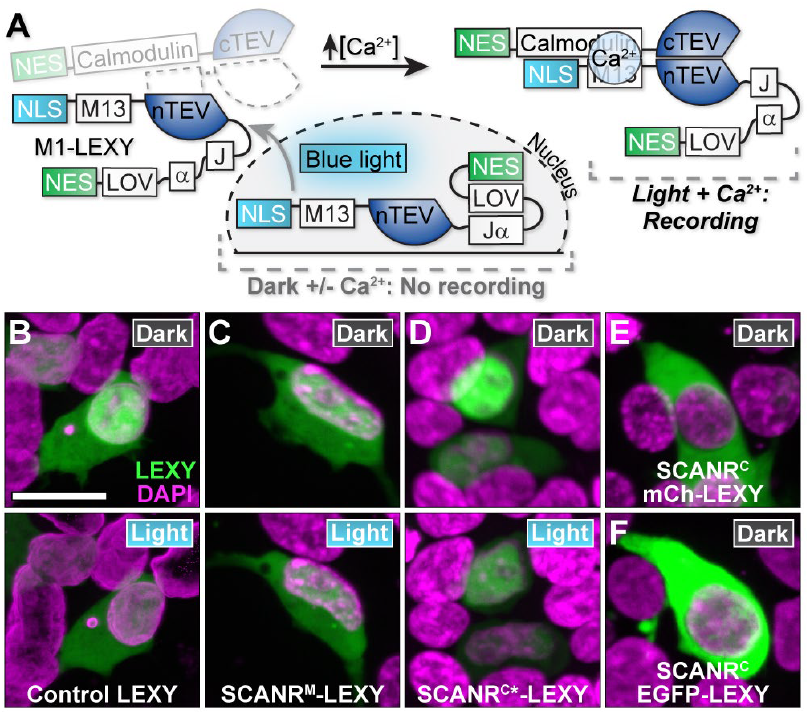
Application of LEXY-based control over subcellular localization to the SCANR system. (**A**) Schematic depiction of the LEXY-modified SCANR constructs. In the dark, the LEXY-modified SCANR construct is designed to be sequestered in the nucleus. Upon the addition of blue light, the LEXY construct is shuttled into the cytoplasm, where, in the presence of high Ca^2+^, it can be reconstituted with the other SCANR half, thus re-forming active TEV protease and turning over substrate. (**B**–**D**) Expression and illumination of LEXY modified constructs. HEK293T cells were transfected with the indicated constructs and illuminated with 488 nm light to induce nucleus-to-cytoplasm translocation. (**B**) The control LEXY plasmid shows clear loss of fluorescence from the nucleus as the protein is shuttled out. (**C**) By visualizing the mCherry component in SCANR^M^-LEXY and (**D**) SCANR^C*^-LEXY we observed a modest nucleus-to-cytoplasm translocation based on the decrease in nuclear signal upon irradiation with blue light. (**E**–**F**) Two other variants of the calmodulin bearing SCANR components, SCANR^C^-mCh-LEXY (*E*) and SCANR^C^-EGFP-LEXY (*F*), localized to the cytoplasm before the addition of blue light, thus prohibiting nucleus-to-cytoplasm-based control of SCANR activation. All images are maximum intensity z-projections of confocal microscopy images. Scale bar = 20 μm.

The LINuS- and LANS-based constructs produced similar challenges for application to SCANR. For example, LINuS hides or exposes a nuclear localization signal to control translocation from the cytoplasm to the nucleus upon exposure to blue light (**Supplementary Figure S1A**). We explored three variants of LINuS (LINuS 9, 10, and 11) in conjunction with M13 SCANR constructs that vary based on the strength of the NLS used in LINuS. In each case with the control constructs, we observed strong localization in the cytoplasm with low levels of nuclear fluorescence in the dark state, and modest translocation to the nucleus upon exposure to blue light (**Supplementary Figure S1B**). However, in the LINuS-modified SCANR constructs, we observed low levels of expression in both the cytoplasm and nucleus (For LINuS 9 and 10 variants) and strong nucleus mis-localization in the dark state for the LINuS 11 variant, precluding the use of these constructs in controlling SCANR activity through limiting access to the nucleus (**Supplementary Figure S1C**). Likewise, the LANS system, which also functions as a blue light activated nuclear importer (**Supplementary Figure S2A**) produced mostly cytoplasmic localization in the dark state and substantial translocation to the nucleus in control, LANS-modified M1 SCANR, and LANS-modified calmodulin SCANR (**Supplementary Figure S2B**). However, the small amount of nuclear LANS-modified SCANR constructs in the dark state would likely contribute to background Ca^2+^ recording. Ultimately, although it may be possible to adapt LEXY, LINuS, and LANS to controlling SCANR Ca^2+^ recording, our experiments with first-generation constructs did not provide confidence that the strategy would be successful, so we explored other options.

### Evaluating application of CRY2-C1B1-based optogenetic control of subcellular localization to SCANR

Next, we explored the application of a popular optogenetics system that functions through protein hetero-dimerization in the presence of blue light (CRY2-C1B1). Essentially, we developed a system to isolate a single half of SCANR in the cytoplasm (C1) while having the other partner (M1) tethered to the cell membrane via a palmitoylation sequence. To accomplish this, we employed the CRY2-C1B1 system, an optogenetics method that shuttles proteins between subcellular compartments based on light-induced heterodimerization^14^. Essentially, CRY2/C1B1 is a heterodimeric protein complex that dimerizes upon activation with blue light. When applied to SCANR, in the presence of blue light, the CRY2-C1 and CRY2-dsRed substrate fusion proteins would be translocated to the nucleus and bind to the C1B1 at the cell membrane (**Figure 3A**). The palmitoylated M1 would be present at the membrane, and, if Ca^2+^ is present, the M1 and C1 constructs would reform the active TEV protease so that it can turn over the dsRed substrate, leading to a fluorescently marked nucleus. Thus, the SCANR system will be active only when both blue light and Ca^2+^ are present.

**Figure 3.**
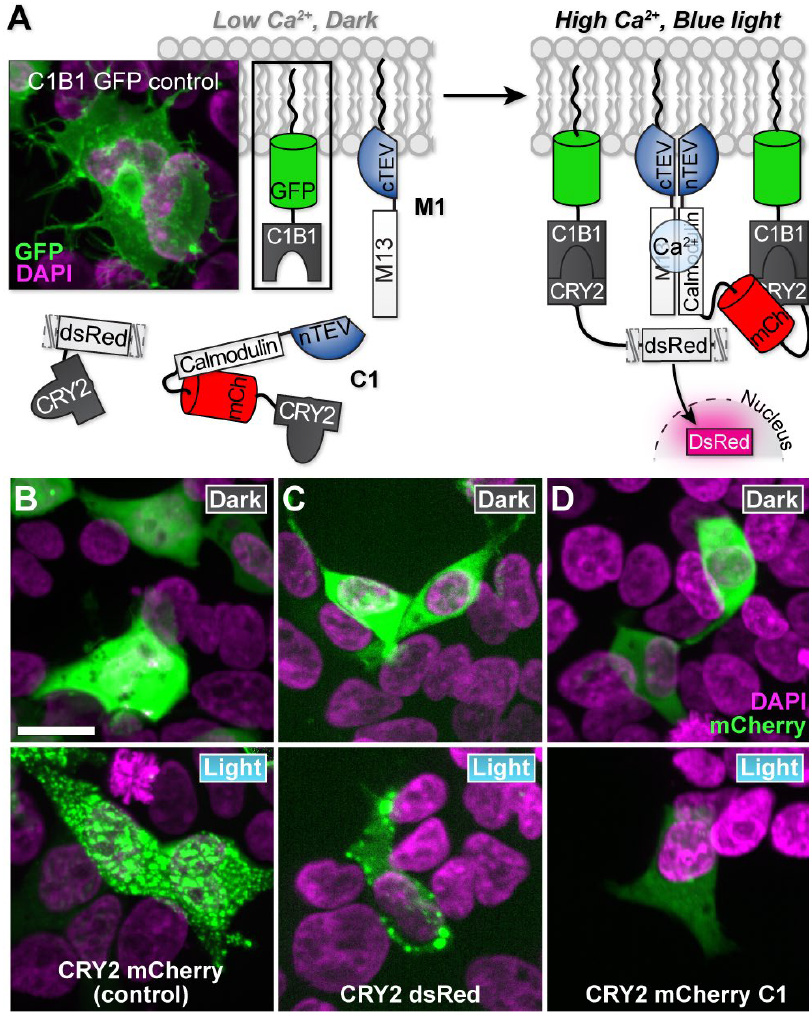
Application of CRY2/C1B1-based cytoplasm-to-plasma membrane translocation for controlling SCANR Ca^2+^ recording. **(A)** Schematic depiction of the strategy for using CRY2/C1B1 for controlling SCANR activity. The CRY2/C1B1-modified SCANR has a palmitoylated M13-bearing SCANR construct (M1) that is tethered to the cell membrane. The calmodulin-bearing SCANR component (C1) and the dsRed substrate are localized to the cytoplasm in the dark state and are designed to be translocated to the cell membrane in the presence of blue light due to the CRY2-C1B1 heterodimerization. Thus, Ca^2+^ recording with this modification of SCANR requires both blue light to induce translocation of all components to the cell membrane, and Ca^2+^ to induce SCANR re-formation. The inset shows a control C1B1-GFP localized to the cell membrane. (**B**–**D**) Evaluation of the expression and light-induced cytoplasm-to-plasma membrane translocation of the CRY2-modified SCANR constructs. HEK293T cells were transfected with the indicated constructs and illuminated with 488 nm light. The Dark (top) and Light (bottom) rows show different locations in the tissue culture plate. They are not images of the same cells at different timepoints. (**B**) The control CRY2-mCherry construct is unmodified from the reported literature construct and shows a distinct cytoplasm-to-membrane distribution upon illumination blue light. (**C**) Like the results with the control construct shown in (*B*), the CRY2-dsRed TEV substrate shown a distinct change in localization upon exposure to blue light. (**D**) Conversely, the CRY2-modified calmodulin component of SCANR (C1) does not show an obvious redistribution from the cytoplasm to the plasma membrane upon exposure to blue light., although there is a distinct loss in fluorescence intensity upon blue light illumination. All images are maximum intensity z-projections of confocal microscopy images. Scale bar = 20 μm.

Using control constructs, we first tested the expression of each construct and confirmed that our optogenetics illumination protocols were able to reliably translocate the separate elements to the cell membrane. Control GFP-labeled C1B1 produced a characteristic membrane pattern of fluorescence (**Figure 3B** and **Supplementary Figure S3**). Likewise, recruitment of its control CRY2 partner upon illumination with blue light revealed membrane labeling analogous to the original literature description of this system (**Figure 3C**). This cytoplasm to cell membrane translocation was less pronounced with the caged dsRED TEV substrate and CRY2-modified calmodulin SCANR (**Figure 3D**).

Ultimately, we found that there was no TEV substrate turnover (fluorescent red nuclei) after recruitment with blue light and an increase in Ca^2+^ concentration with ionomycin. It has been shown by others that this protein-protein interaction is not as successful when C1B1 is tethered to the cell membrane, as it is utilized in our system^15^. We hypothesize that this lack of output may also be attributed to specific protein conformations that are required for activity, which may be more challenging to adopt due to the topological constraints imposed by membrane localization.

### Activating SCANR using a chemogenetic strategy for controlling a protein’s subcellular localization

Our final attempt to control SCANR recording though manipulating its component’s subcellular localization employed a small-molecule-controlled protein targeting system developed by Crabtree and coworkers^16^. In this system, three FKBPs (FKBP3x) are linked to a protein of interest downstream of a nuclear localization sequence. The FRB, in the presence of rapamycin, will retrieve the FKBP construct out of the nucleus and bring it into the cytoplasm (**Figure 4A**). We created a fusion protein composed of the C-terminal SCANR component, a nuclear localization sequence, and three sequential FKBPs (C1-NLS-FKBP3x). This construct is designed to reside in the nucleus where it is segregated from the N-terminal SCANR construct, which is trapped in the cytoplasm by the addition of an ERT2 domain. Thus, in this system, SCANR activation and turnover of dsRed substrate to produce a fluorescent nucleus would only occur in the presence of both high Ca^2+^ and rapamycin-induced translocation of C1-NLS-FKBP3x to the cytoplasm.

**Figure 4.**
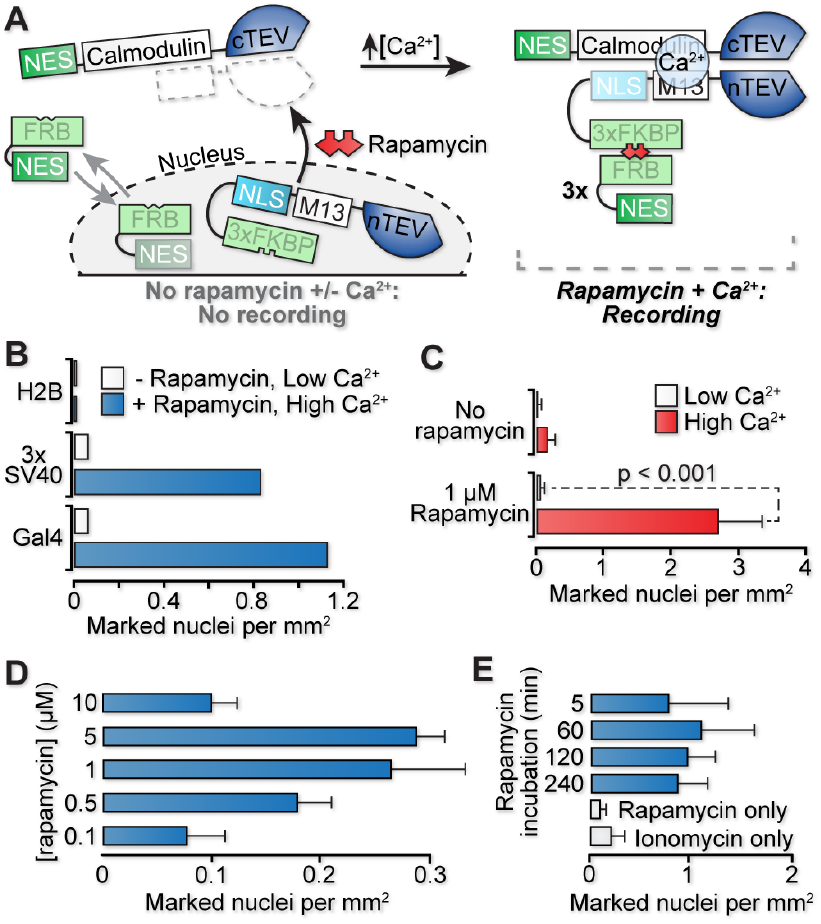
Engineering chemogenetic control of SCANR Ca^2+^ recording using rapamycin-induced heterodimerization. (**A**) Schematic of the rapamycin-inducible SCANR. The modified SCANR system utilizes an FRB with a weak NES that is in equilibrium between the cytoplasm and nucleus. Upon the addition of rapamycin, the three FRB-NES units dimerizes with a nuclear 3xFKBP-SCANR-NLS, overpowering the NLS and pulling the protein complex into the cytoplasm. The FKBP-SCANR construct can then interact with its SCANR counterpart when induced to do so by the high Ca^2+^, resulting in fluorogenic substrate turnover. (**B**) Comparison of nuclear localization signals in the rapamycin-inducible SCANR in transiently transfected HEK293T cells. The Gal4, as an NLS, had the greatest signal to noise ratio, whereas the H2B protein was a poor NLS with no signal even in the presence of both high Ca^2+^ and rapamycin. (**C**) Using the Gal4 as the NLS, rapamycin-inducible SCANR was transiently transfected into HEK293T cells and the cells were treated +/-rapamycin, and +/-ionomycin (to induce high Ca^2+^). Analysis of substrate turnover based on the presence of fluorescently marked nuclei shown a dependence on exposure to both rapamycin and high Ca^2+^. (D) Increase in rapamycin-inducible SCANR marked nuclei with increasing rapamycin concentration. (**E**) Rapamycin-inducible SCANR constructs expressed in HEK293T cells were exposed to 1 μM rapamycin at varying times prior to ionomycin-induced high Ca^2+^. Incubation with rapamycin for as little as 5 min prior to high Ca^2+^ resulted in the same level of signal as a 240 min rapamycin exposure. Error bars represent standard deviation for sample sizes of six (*C*), and three (*D, E*).

We began our exploration of the rapamycin inducible system by identifying the optimal nuclear localization signal for sequestering the SCANR components in the nucleus sans rapamycin and promoting nuclear export and SCANR recording in the presence of rapamycin and Ca^2+^. As candidate nuclear localization signals, we chose Gal4-VP16, 3x-NLS from SV40, and the entire H2B protein. We cloned each construct, expressed them with the appropriate SCANR counterpart and a profluorescent TEV substrate, and assayed for substrate turnover in the presence and absence of ionomycin-induced high Ca^2+^ concentrations in HEK293T cells. Interestingly, these experiments revealed a degree of rapamycin-induced control of SCANR recording in each instance, although the best performing nuclear localization signal was the Gal4-VP16, which had the greatest increase in signal with essentially no background in the absence of the ionomycin-induced Ca^2+^ spike (**Figure 4B**). Importantly, using the Gal4 NLS, rapamycin-inducible SCANR showed no significant substrate turnover without the addition of rapamycin or the induction of Ca^2+^ concentration spikes (**Figure 4C**). Calcium recording with SCANR in this system required both high Ca^2+^ and an ionomycin stimulus, thus permitting control of the SCANR recording time through the addition of rapamycin.

Having identified a suitable NLS and construct pair for controlling SCANR with rapamycin, we began characterizing dose response with respect to rapamycin incubation time and concentration. We found that the SCANR signal increased with increasing ionomycin concentration until we approached the rapamycin solubility limit at 5 μM, at which point the signal number decreased, most likely due to rapamycin solubility in water decreasing at around 10 μM (**Figure 4D**).

Another important consideration is the required rapamycin exposure time to achieve Ca^2+^ recording. To asses this parameter, we exposed SCANR transfected HEK293T cells to rapamycin for 5 min to 4 h before treatment with ionomycin. Analysis of substrate turnover as indicated by marked nuclei revealed significant numbers of marked nuclei after only 5 min, and no increase in recorded nuclei with longer incubation times up to 4 h, indicating that rapamycin binding to the SCANR component and translocation out of the nucleus occurs on the minute time scale (**Figure 4E**).

Ultimately, this modification was straightforward to implement from a cloning standpoint and produced robust control of SCANR recording on the minute timescale. This strategy provides an avenue for controlling the onset of Ca^2+^ recording in situations where light application and optogenetics are not applicable, such as over long distances in thick organisms or in organisms and behavioral systems incompatible with optogenetics setups.

### Applying LOV-Jα-based steric control of protein association to the SCANR system

Finally, we evaluated utilizing the LOV-Jα optogenetic steric block to directly cage the SCANR components. The LOV-Jα system has been extensively used to sterically block protein-protein interactions in mammalian cells. As applied to SCANR, the LOV-Jα domain would be engineered to block the interaction of the two separate SCANR components, either from the Ca^2+^ binding site or protein surface needed for TEV reconstitution (**Figure 5A**). To begin, we produced constructs with the LOV-Jα domain positioned at the amino terminus of either the calmodulin or M13-containing component of SCANR. We confirmed the expression of these constructs in HEK293T cells, revealing strong expression of the LOV-Jα modified calmodulin SCANR construct and only meager expression of the M13 SCANR variant (**Supplementary Figure S4**).

**Figure 5.**
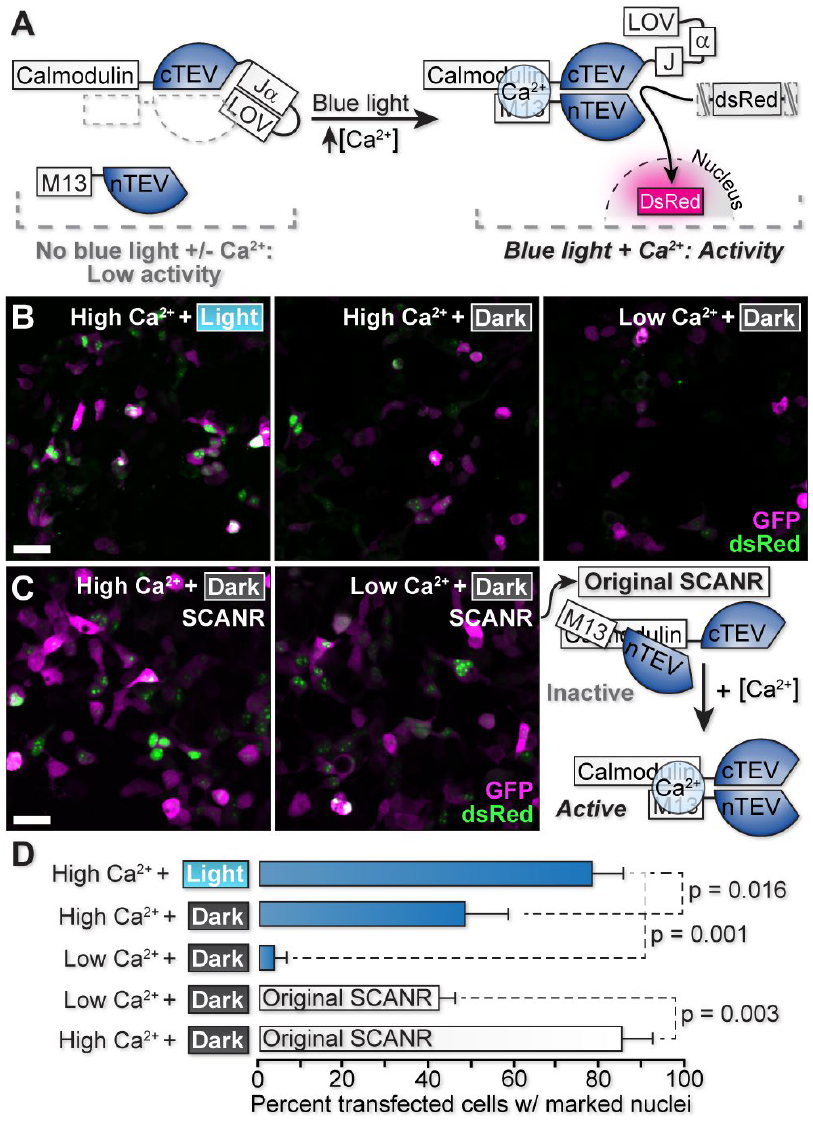
Characterization and comparison of LOV-modified and original SCANR. **(A)** Schematic of the LOV-modified SCANR system. The LOV-linked SCANR component is designed to be sterically hindered from associating with its SCANR counterpart and cleaving substrate in the dark state. Only upon the addition of blue light and high Ca^2+^ can LOV-modified SCANR be dimerize to become active and cleave substrate. (**B**) Expression and illumination of LOV-modified SCANR, which shows similar transfection efficiencies (GFP) with the dsRed nuclear labelling being most prominent when cells are exposed to both blue light and high Ca^2+^. (**C**) In comparison, unmodified SCANR shows much higher background of dsRed labelling in the absence of high Ca^2+^, and similar levels of mark nuclei compared to the LOV-modified SCANR exposed to both blue light and Ca^2+^. (**D**) Quantification of SCANR marked nuclei from the experiments shown in (*B*) and (*C*). All images are maximum intensity z-projections of confocal microscopy images. Error bars represent standard deviation with a sample size of 3. Scale bar = 20 μm.

Next, we evaluated the light dependence of Ca^2+^ recording with the calmodulin SCANR construct. As noted previously, the original SCANR constructs produced background Ca^2+^ recording signal in HEK293 cells that were not exposed to the ionophore ionomycin, due to the occasional naturally-occurring Ca^2+^ concentration spikes that occur in this cell line (*e*.*g*., in response to media changes^17^). Importantly, with the LOV-Jα modified calmodulin SCANR, we observed essentially no substrate turnover, indicating that this modified SCANR was less likely to reconstitute active enzyme in the absence of blue light exposure. We then explored substrate turnover upon exposure to ionomycin with three exterior Ca^2+^ buffer concentrations, with and without blue light exposure. (**Supplementary Figure S5**).

When no Ca^2+^ was added to the external buffer, so that Ca^2+^ concentration increase by ionomycin exposure relies on internal stores of Ca^2+^, we saw an increase in SCANR marked nuclei upon exposure to ionomycin that was not statistically significant. Increasing the buffer concentration to 315 μM produced a more robust and statistically significant increase in SCANR marked nuclei. Further increases in the external Ca^2+^ supply were detrimental to substrate turnover, which could be a reflection of the complex dynamics involved with the ionomycin induced release of Ca^2+^. For example, there are two phases to the mechanism of induced Ca^2+^ flux via ionomycin: (1) release of Ca^2+^ from intracellular stores and (2) crossing of Ca^2+^ from the extracellular environment^18^. This is a complex process that in certain conditions leads to cell death, which may bias results in the very high Ca^2+^ situations.

Finally, we generated a construct bearing the LOV-Jα modified calmodulin SCANR, the original M13-modified SCANR, and an intervening GFP for marking expressing cells, in a multicistronic p2A vector. Expression of this construct and the dsRed pro-fluorescent TEV protease substrate in HEK293 cells revealed very low levels of marked nuclei in dark, low Ca^2+^ situations, a modest 50% of transfected cells in high Ca^2+^ and in the dark, and a more robust ~80% of transfected cells marked when high Ca^2+^ and blue light were applied (**Figure 5B, D**). In contrast, the original SCANR constructs in this assay produced much higher background in the absence of ionomycin-induced Ca^2+^ concentration spikes, rivaling the signal observed in high Ca^2+^ LOV-modified SCANR in dark conditions (**Figure 5C, D**). Importantly, the original SCANR + Ca^2+^ had levels of fluorophore-marked nuclei similar to the LOV-modified SCANR + Ca^2+^ and blue light, suggesting that the addition of the LOV domain does not drastically decrease the upper limits on the construct’s activity.

## CONCLUSIONS

In summary, we have shown that the original SCANR constructs are amenable to several modifications that permit direct steric or subcellular localization-based control of activity. Through testing of different commercially available options, both the LOV-Jα constructs and the FKBP/FRB constructs were successful in reducing background signal and only showed significant turn-on in the presence of both increased Ca^2+^ and either blue light or rapamycin, respectively. To increase the utility of this system, more diverse protease substrates such as zipGFP^19^ or quenched GFP with hydrophobic peptides^20,21^, could be utilized to better understand SCANR kinetics and produce a faster turn on of signal. Additionally, application of photo-activatable rapamycin, which has been recently applied to controlling split TEV activation^22^, would provide an additional level of control over this system. Future work will be dedicated to showing the utility of modified SCANR in neurons by showing reduced background due to spontaneous action potentials. Once the cultured primary neuron work has been validated, we will test our system in zebrafish to show the marking of permanent Ca^2+^ signaling in the brain in response to various stimuli.

## ASSOCIATED CONTENT

### Supporting Information

The Supporting Information is available free of charge on the ACS Publications website.

General materials and methods and specific procedures for the cloning of all constructs, cell transfection protocols, ionomycin exposure assays, fluorescence microscopy methods and image quantification details, optogenetics illumination methods and rapamycin exposure procedures, supplementary figures and cloning primer tables. (PDF).

## AUTHOR INFORMATION

### Author Contributions

S.T.L. conceived the project and wrote the manuscript. Y.Z. and B.K.C. performed experiments and data analysis and wrote the manuscript, A.S. and K.E. performed experiments and data analysis.

### Notes

The authors declare no competing financial interests.

## ACKNOWLEDGMENT

We thank A. Guo and M. Shelly for helpful discussions and experimental expertise, A. Preston, D. Cervasio, and P. Kumar for critical reading of the manuscript, and the Stony Brook University Genomics Center.

## ABBREVIATIONS

CAMPARI: Calcium modulated Photoactivatable Ratiometric Integrator
CRY2: Cryptochrome circadian regulator 2
DMEM: Dulbecco’s Modified Eagle Medium
FKBP: FK506 binding protein
FLARE: Fast Light- and Activity-Regulated Expression
FRB: FKBP-rapamycin binding protein
HBSS: Hank’s Balanced Salt Solution
LANS: Light Activated Nuclear Shuttle
LEXY: Light-inducible nuclear export system
LINuS: Light-inducible Nuclear Localization Signal
LOV: Light Oxygen Voltage
NLS: Nuclear Localization Signal
PBS: Phosphate Buffered Saline
ROI: Region of Interest
SCANR: Split Tobacco Etch Virus (TEV) protease Calcium-regulated Neuron Recorder
TRIC: Transcriptional reporter of Intracellular Ca^2+^.

## REFERENCES

(1) Chen, T.-W., Wardill, T. J., Sun, Y., Pulver, S. R., Renninger, S. L., Baohan, A., Schreiter, E. R., Kerr, R. A., Orger, M. B., Jayaraman, V., Looger, L. L., Svoboda, K., and Kim, D. S. (2013) Ultrasensitive fluorescent proteins for imaging neuronal activity. Nature 499, 295–300.

(2) Akerboom, J., Rivera, J. D. V., Guilbe, M. M. R., Malavé, E. C. A., Hernandez, H. H., Tian, L., Hires, S. A., Marvin, J. S., Looger, L. L., and Schreiter, E. R. (2009) Crystal structures of the GCaMP calcium sensor reveal the mechanism of fluorescence signal change and aid rational design. J. Biol. Chem. 284, 6455–64.

(3) Tian, L., Hires, S. A., Mao, T., Huber, D., Chiappe, M. E., Chalasani, S. H., Petreanu, L., Akerboom, J., McKinney, S. A., Schreiter, E. R., Bargmann, C. I., Jayaraman, V., Svoboda, K., and Looger, L. L. (2009) Imaging neural activity in worms, flies and mice with improved GCaMP calcium indicators. Nat. Methods 6, 875–881.

(4) Muto, A., and Kawakami, K. (2011) Imaging functional neural circuits in zebrafish with a new GCaMP and the Gal4FF-UAS system. Commun. Integr. Biol. 4, 566–8.

(5) Fosque, B. F., Sun, Y., Dana, H., Yang, C.-T., Ohyama, T., Tadross, M. R., Patel, R., Zlatic, M., Kim, D. S., Ahrens, M. B., Jayaraman, V., Looger, L. L., and Schreiter, E. R. (2015) Labeling of active neural circuits in vivo with designed calcium integrators. Science (80-.). 347, 755–60.

(6) Gao, X. J., Riabinina, O., Li, J., Potter, C. J., Clandinin, T. R., and Luo, L. (2015) A transcriptional reporter of intracellular Ca2+ in Drosophila. Nat. Neurosci. 18, 917– 925.

(7) Wang, W., Wildes, C. P., Pattarabanjird, T., Sanchez, M. I., Glober, G. F., Matthews, G. A., Tye, K. M., and Ting, A. Y. (2017) A light-and calcium-gated transcription factor for imaging and manipulating activated neurons. Nat. Biotechnol. 35, 864–871.

(8) O’Neill, B. K., and Laughlin, S. T. (2018) Neuronal Calcium Recording with an Engineered TEV Protease. ACS Chem. Biol. 13, 1159–1164.

(9) Wehr, M. C., Laage, R., Bolz, U., Fischer, T. M., Grünewald, S., Scheek, S., Bach, A., Nave, K.-A., and Rossner, M. J. (2006) Monitoring regulated protein-protein interactions using split TEV. Nat. Methods 3, 985– 93.

(10) Niopek, D., Wehler, P., Roensch, J., Eils, R., and Di Ventura, B. (2016) Optogenetic control of nuclear protein export. Nat. Commun. 7, 10624.

(11) Yumerefendi, H., Dickinson, D. J., Wang, H., Zimmerman, S. P., Bear, J. E., Goldstein, B., Hahn, K., and Kuhlman, B. (2015) Control of Protein Activity and Cell Fate Specification via Light-Mediated Nuclear Translocation. PLoS One 10, e0128443.

(12) Wu, Y. I., Frey, D., Lungu, O. I., Jaehrig, A., Schlichting, I., Kuhlman, B., and Hahn, K. M. (2009) A genetically encoded photoactivatable Rac controls the motility of living cells. Nature 461, 104–8.

(13) Niopek, D., Benzinger, D., Roensch, J., Draebing, T., Wehler, P., Eils, R., and Di Ventura, B. (2014) Engineering light-inducible nuclear localization signals for precise spatiotemporal control of protein dynamics in living cells. Nat. Commun. 5, 4404.

(14) Kennedy, M. J., Hughes, R. M., Peteya, L. A., Schwartz, J.., Ehlers, M. D., and Tucker, C. L. (2010) Rapid blue-light-mediated induction of protein interactions in living cells. Nat. Methods 7, 973–5.

(15) Benedetti, L., Barentine, A. E. S., Messa, M., Wheeler, H., Bewersdorf, J., and De Camilli, P. (2018) Light-activated protein interaction with high spatial subcellular confinement. Proc. Natl. Acad. Sci. 115, E2238–E2245.

(16) Klemm, J. D., Beals, C. R., and Crabtree, G. R. (1997) Rapid targeting of nuclear proteins to the cytoplasm. Curr. Biol. 7, 638–44.

(17) Tong, J., Du, G. G., Chen, S. R. W., and Maclennan, D. H. (1999) HEK-293 cells possess a carbachol-and thapsigargin-sensitive intracellular Ca2+ store that is responsive to stop-flow medium changes and insensitive to caffeine and ryanodine. Biochem. J. 343.

(18) Morgan, A. J., and Jacob, R. (1994) Ionomycin enhances Ca2+ influx by stimulating store-regulated cation entry and not by a direct action at the plasma membrane. Biochem. J. 300 (Pt 3), 665–72.

(19) To, T.-L., Schepis, A., Ruiz-González, R., Zhang, Q., Yu, D., Dong, Z., Coughlin, S. R., and Shu, X. (2016) Rational Design of a GFP-Based Fluorogenic Caspase Reporter for Imaging Apoptosis In Vivo. Cell Chem. Biol. 23, 875–882.

(20) Nicholls, S. B., Chu, J., Abbruzzese, G., Tremblay, K. D., and Hardy, J. A. (2011) Mechanism of a Genetically Encoded Dark-to-Bright Reporter for Caspase Activity. J. Biol. Chem. 286, 24977–24986.

(21) Nicholls, S. B., and Hardy, J. A. (2013) Structural basis of fluorescence quenching in caspase activatable-GFP. Protein Sci. 22, 247–257.

(22) Caldwell, R. M., Bermudez, J. G., Thai, D., Aonbangkhen, C., Schuster, B. S., Courtney, T., Deiters, A., Hammer, D. A., Chenoweth, D. M., and Good, M. C. (2018) Optochemical Control of Protein Localization and Activity within Cell-like Compartments. Biochemistry 57, 2590–2596.

