## Supporting Information for "Chemogenetic and optogenetic strategies for spatiotemporal control of split-enzyme-based calcium recording"

### General materials and methods

The fidelity of all DNA constructs was confirmed by DNA sequencing at the Stony Brook University DNA Sequencing facility. Mouse α-myc tag IgG1 (Clone Myc.A7) was purchased from Abcam (Cambridge, United Kingdom). Oligonucleotides were purchased from Integrated DNA Technologies (Coralville, IA). TOP10 One-Shot and Mach1 competent cells, Lipofectamine 3000, and cell culture media were purchased from Invitrogen (Carlsbad, CA). Secondary antibodies Alexa Fluor® 647-conjugated AffiniPure goat α-mouse IgG (H+L) and Alexa Fluor® 488 conjugated AffiniPure goat α-rabbit IgG (H+L) were purchased from Jackson ImmunoResearch Laboratories, Inc. (West Grove, PA). Phusion DNA polymerase, Taq DNA polymerase, calf intestinal alkaline phosphatase (CIP), and restriction enzymes were purchased from New England BioLabs (Beverly, MA). Omega gel extraction kit and Omega E.Z.N.A. plasmid mini kit II was purchased from Omega Bio-Tek, Inc. (Norcross, GA). Ionomycin was purchased from Santa Cruz Biotechnology, Inc. (Dallas, TX). Vectashield with DAPI was purchased from VWR (Radnor, PA). Thermo Scientific Nunc Lab-Tek II Chamber slides (8 well) were purchased from Fisher Scientific (Boston, MA).

### Origin of DNA PCR templates and plasmids used in construction of SCANR constructs

SYS-V1046 (dsRed substrate plasmid) was purchased from Systasy Bioscience (Munich, Germany).[1] DN41, pDN42, pDN43, and NLS-mCherry-LEXY (pDN122 were a gift from Barbara Di Ventura & Roland Elis (Addgene plasmid # 61343, 61344, 61345, 72655). pTriEX-mCherry::LANS1 and pTriEX-mCherry::LANS4 were a gift from Brian Kuhlman (Addgene plasmid # 60784, 60785). pCRY2PHR-mCherryN1 and pCIBN(deltaNLS)-pnGFP were a gift from Chandra Tucker (Addgene plasmid # 26866, 26867). 3xFKBP-myc and FRB-myc were a gift from Carolyn Bertozzi (Addgene plasmid # 20213, 20228). pGP-CMV-NES-jRGECO1a was a gift from Douglas Kim (Addgene plasmid # 61563). H2B-mCherry was a gift from Robert Benezra (Addgene plasmid # 20972).

### Cloning of modified SCANR and control expression vectors

With the exception of CRY2 constructs, M1-3xFKBP, C1 Gal4 VP16 FKBP3x, and C1-3xNLS FKBP3x, constructs were prepared using the Circular Polymerase Extension Cloning (CPEC) method, which uses overlapping regions in the DNA to assemble a linear vector component and all required inserts into a circular plasmid product, as described previously.[2] The complete list of primers is detailed in Tables S1–S5.

The SCANR constructs for CRY2-C1B1 assays were built into the pCRY2PHR vector (Addgene #26866). The palmitoylated SCANR constructs were built into the pCIBN(deltaNLS)-pnGFP vector (Addgene #26867). The SCANR constructs for LEXY assays were built into the NLS-mCherry-LEXY vector (Addgene #72655). The SCANR constructs for LINUS and LANS assays were built into the pDN41-43 vectors (Addgene #'s 61343, 61344, 61345) and pTriEX-mCherry::LANS1 and 4 vectors respectively (Addgene #'s 60784, 60785). The SCANR constructs for rapamycin assays were built into the 3xFKBP-myc vector (Addgene #20213). The SCANR constructs for LOV assays were built from the PA-Rac1 vector into our constructed SCANR constructs.[3]

The large linear vector components for CPEC were amplified by PCR using a thermocycler program with 30 cycles of 98 °C for 30 s, 54 °C for 30 s, and 72 °C for 2–6min. For PCR of the smaller CPEC reaction inserts, the thermocycler program was 30 cycles of 98 °C for 30 s, 54 °C for 30 s, and 72°C for 30 s. The PCR products for both the linear vectors and inserts were purified prior to use in CPEC reactions by agarose gel electrophoresis, product excision from the gel, and product isolation using the Omega gel extraction kit. The linker region in the LEXY constructs (Table S1) was constructed by slowly annealing the two primers together from 98°C to 60°C over the course of an hour, then allowed to cool to room temperature.

To combine the vector and inserts by CPEC, a two-step CPEC reaction was performed. First, each individual insert required for a particular construct was added to the CPEC reaction at a 1:1 molar ratio and the inserts were combined into a single unit by thermocycling for 25 cycles of 98 °C for 10 s, slow ramp annealing from 70 to 55 °C (1 °C/s), 55 °C for 30 s, and 72 °C for 30 s. After this first reaction was complete, the linear vector was added into the reaction at molar ratio of 1:2 (vector:insert) and the target plasmid was assembled by thermocycling for 25 cycles of 98 °C for 10 s, slow ramp anneal from 70 to 55 °C (1 °C/s), 55 °C for 30 s, and 72 °C for 2 min. In situations where the target DNA construct required the addition of only one insert, a single CPEC reaction using the molar ratios and thermocycler program for the second step described above was done. Finally, 4 µL of the CPEC reaction was transformed into Mach1 or TOP10 One-Shot chemically competent cells and individual clones were isolated and analyzed for proper insert incorporation by DNA sequencing.

The expression vector for C1, M1, and dsRed substrate into the CRY2 vector (Addgene plasmid pCRY2PHR) was constructed using traditional restriction digest cloning using XmaI and NotI restriction sites, which are included in the PCR primer sequences (Table S2). H2B into the M1-3xFKBP vector was constructed using traditional restriction digest cloning using XbaI and XhoI restriction sites, which are included in the PCR primer sequences (Table S4). Gal4-VP16 and 3xNLS into the C1-3xFKBP vector was constructed using traditional restriction digest cloning using HindIII and XbaI restriction sites, which are included in the PCR primer sequences (Table S4). Essentially, the desired insert was amplified using PCR with thermocycling at 30 cycles of 98 °C for 30 s, 54 °C for 30 s, and 72°C for 30 s, and the product was purified by excision from an agarose gel after electrophoresis and isolation using the Omega gel extraction kit. The destination vector was digested with the appropriate enzymes in the presence of CIP and purified by excision from an agarose gel followed by gel extraction using the Omega gel extraction kit before ligation at room temperature for 10 minutes with T4 DNA ligase. Finally, the plasmid was transformed into Mach1 chemically competent cells and individual clones were isolated and analyzed for proper insert incorporation by DNA sequencing.

Mutagenesis of M1 H2B FKBP3x (Table S4) was produced using the linearization PCR primers. The linear vector was amplified by PCR using a thermocycler program with 30 cycles of 98 °C for 30 s, 54 °C for 30 s, and 72 °C for 2–6min. The band was purified by agarose gel electrophoresis, product excision from the gel, and product isolation using the Omega gel extraction kit. The linear mutated vector was circularized without nicks via thermocycling for 25 cycles of 98 °C for 10 s, slow ramp annealing from 70 to 55 °C (1 °C/s), 55 °C for 30 s, and 72 °C for 30 s. Finally, 4 μL of the CPEC reaction was transformed into Mach1 or TOP10 One-Shot chemically competent cells and individual clones were isolated and analyzed for proper mutation by DNA sequencing.

### HEK293T cell culture and transfection

HEK293T cells were maintained in high-glucose Dulbecco's Modified Eagle Medium (DMEM) supplemented with 5% FBS and 50,000 U penicillin-streptomycin at 37 °C in air with 5% CO2. Cells were passaged on alternating days when they reached approximately 85% confluency. Cells were transfected at 75–85% confluence in Millipore 8-chamber culture slides or 35 mm culture dishes with Lipofectamine 3000 according to the manufacturer's instructions. Cells were cultured for 24 h before ionomycin assays or immunohistochemistry analyses were performed.

### Ionomycin assay for increasing intracellular $Ca^{2+}$ in HEK293T cells

Assay was performed as described previously[3]. Briefly, media was removed from the cells and the cells were washed twice with 400 μL phosphate buffered saline (PBS) pH 7.4. Cells were incubated in 5 μM ionomycin in Hanks balanced salt solution (HBSS) at room temperature. The ionomycin solution was removed via aspiration and the cells were washed 2 times with 400 μL PBS to remove all traces of ionomycin. The cells were put into fresh DMEM solution and allowed to incubate for 24 h at 37 °C in air with 5% $CO_2$. Cells in 35 mm plates were put into carbon dioxide independent media and imaged directly by confocal microscopy. Manual cell counting was preformed using a Zeiss Axio Vert A. 1 inverted microscope equipped with a Zeiss EC Plan-Neofluor 20x NA0.50 objective.

### Quantification of DsRed-positive nuclei in SCANR-expressing HEK293T cells ± ionomycin

24 h past exposure of modified SCANR and caged DsRed substrate- expressing HEK293T cells to ionomycin ± blue light or rapamycin, the number of DsRed-fluoresenct nuclei per field of view of the live cells was manually counted with a Zeiss Axio Vert A.1 inverted microscope equipped with a Zeiss EC Plan-Neofluar 20x NA0.50 objective.

### Immunohistochemistry analysis of HEK293 cells

Transfected cells were washed with 400 μL PBS at 24 h post transfection and incubated in 4% paraformaldehyde in PBS for 15 minutes at room temperature. The cells were washed and then permeabilized for 3 min at room temperature by treatment with a solution of 1% bovine serum albumin (BSA) and 0.1% Triton-X 100 in PBS. After washing three times with 400 μL of PBS, the cells were blocked with 1% BSA in PBS for 10 minutes. For cells being probed with α-myc, cells were incubated 1:200 α-myc overnight at 4 °C in 1% BSA in PBS. The cells were washed 3 times with 400 μL PBS before incubation with a 1:100 dilution of goat α-mouse Alexa Flour® 647 in 1% BSA/PBS overnight at 4 °C. Subsequently, the cells were washed 3 times with 400 μL PBS before addition of vectashield with DAPI was added to the plate before addition of the coverslip and subsequent imaging with confocal microscopy.

### Blue light assays for optogenetic strategies

Blue light stimulation for CRY2/C1B1was performed on live cells using a Zeiss Axio Vert A. 1 inverted microscope equipped with a Zeiss EC Plan-Neofluor 20x NA0.50 objective. The green channel excitation is 480/40. Cells were illuminated with blue light while looking through the eye-pieces for no more than 3 s at a time per field of view. The illumination was performed before, and halfway through the ionomycin incubation.

Blue light stimulation for the LOV constructs was performed on live cells using a Zeiss Axio Examiner.D1 modified with an Andor Differential Scanning Disk 2 confocal unit equipped with a 10x NA0.3 air objective and Piezo objective holder for acquiring Z-stacks. Optimal z-stack step size was calculated using the Andor iQ3 software to provide Nyquist sampling. Excitation filter used was: Blue channel excitation 390/40 at 75% intensity at 200 ms for 12 repeats (30 s intervals) for 6 min.

Blue light stimulation for the LEXY, LINUS, and LANS constructs were performed on live cells using a Zeiss Axio Examiner.D1 modified with an Andor Differential Scanning Disk 2 confocal unit equipped with a 40x NA1.0 water immersion objective and Piezo objective holder for acquiring Z-stacks. Optimal z-stack step size was calculated using the Andor iQ3 software to provide Nyquist sampling. Excitation filter used was: Blue channel excitation 390/40 at 100% intensity at (LEXY) 40 ms exposure for 80 repeats (30 s intervals) for 40 min, (LINUS) 1 s exposure for 30 repeats (30 s intervals) for 15 min, and (LANS) 300 ms exposure for 300 repeats (5 s intervals) for 25 min.

### Rapamycin assays for FKBP/FRB dimerization

Media was removed from cells, and rapamycin added at the time specified (0 min to 24 h) before the 5 min ionomycin bathing. Cells were incubated with indicated rapamycin concentration (100 nM-10 µM rapamycin) in DMEM and allowed to incubate for the described time before addition of 5 µM ionomycin/rapamycin/HBSS. After the ionomycin assay, cells were washed with 400 µL PBS, pH 7.4 cells then were placed into rapamycin/DMEM and allowed to incubate for 24 h at 37 °C in air with 5% CO2.

For inhibition with ascomycin, cells were incubated in 100 nM rapamycin and 10 uM ascomycin and treated similarly to the control assays. After the ionomycin assay, cells were washed with 400 µL PBS, pH 7.4, before being placed into 10 uM ascomycin/DMEM, at the indicated time post ionomycin, overnight at 37 °C in air with 5% CO2. Manual cell counting was preformed using a Zeiss Axio Vert A. 1 inverted microscope equipped with a Zeiss EC Plan-Neofluor 20x NA0.50 objective.

### Confocal imaging and image analysis

Confocal microscopy was performed using a Zeiss Axio Examiner.D1 modified with an Andor Differential Scanning Disk 2 confocal unit equipped with a 40x NA1.0 water immersion objective and Piezo objective holder for acquiring Z-stacks. Optimal z-stack step size was calculated using the Andor iQ3 software to provide Nyquist sampling. Excitation and emission filters used were: Blue channel excitation 390/40, emission 452/45; Green channel excitation 482/18, emission 525/45; Red channel excitation 556/20, emission 609/54; Far red channel excitation 640/14. Emission 676/29. Confocal images were converted to maximum intensity projections of the z-stack and level-normalized across all images using ImageJ. Image cropping and organization into figure illustrations was performed using Adobe Illustrator CC. Fixed cells were randomly sampled for each condition. A maximum intensity z-projection of the red channel confocal image for each cell was created containing the slices with the nucleus. An ROI was drawn around the nucleus of each cell and the mean intensity of the pixels in that region was collected.

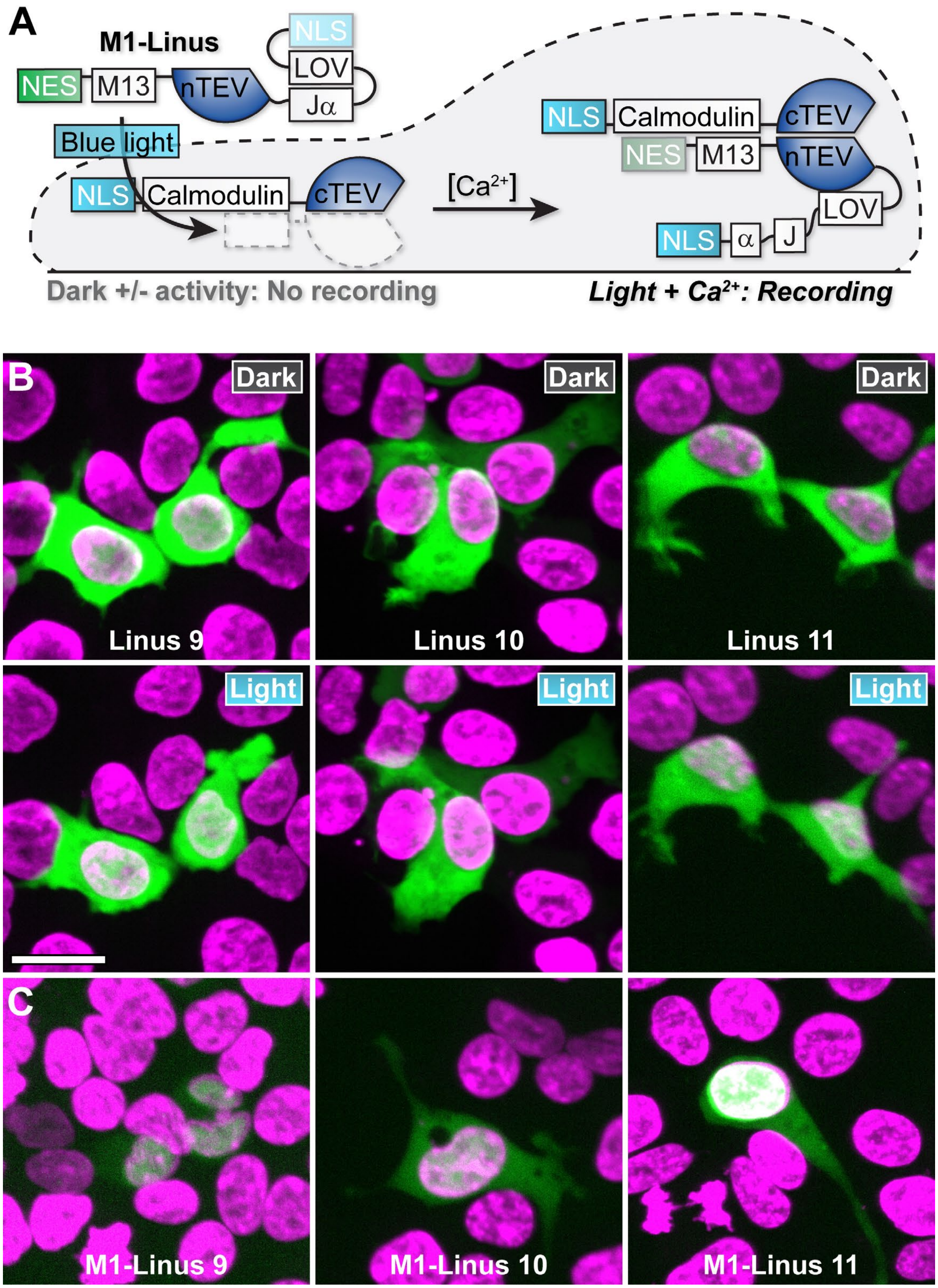

**Figure S1: Application of LiNUS-based control over subcellular localization to the SCANR system.** (**A**) Schematic depiction of the LiNUS modified SCANR constructs. In the dark, the LiNUS-modified SCANR construct is designed to be secluded from the nucleus and localized in the cytoplasm. Only in the presence of blue light can the construct be shuttled into the nucleus, where, in the presence of high $Ca^{2+}$, it can interact with its partner to reforme an active TEV protease. (**B**) Expression and illumination of LiNUS controls. HEK293T cells were transfected with the indicated constructs and illuminated with 488 nm light to induce cytoplasm-to-nucleus translocation. The control LiNUS plasmids showed clear increase of nuclear fluorescence, indicating that the protein was shuttled into the nucleus. (**C**) Visualization of LiNUS-modified M13-containing SCANR localization when expressed in HEK293T cells shows that the dark state protein localization is predominantly in the nucleus. Images are maximum intensity z-projections of confocal microscopy images. Scale bar = 20 µm

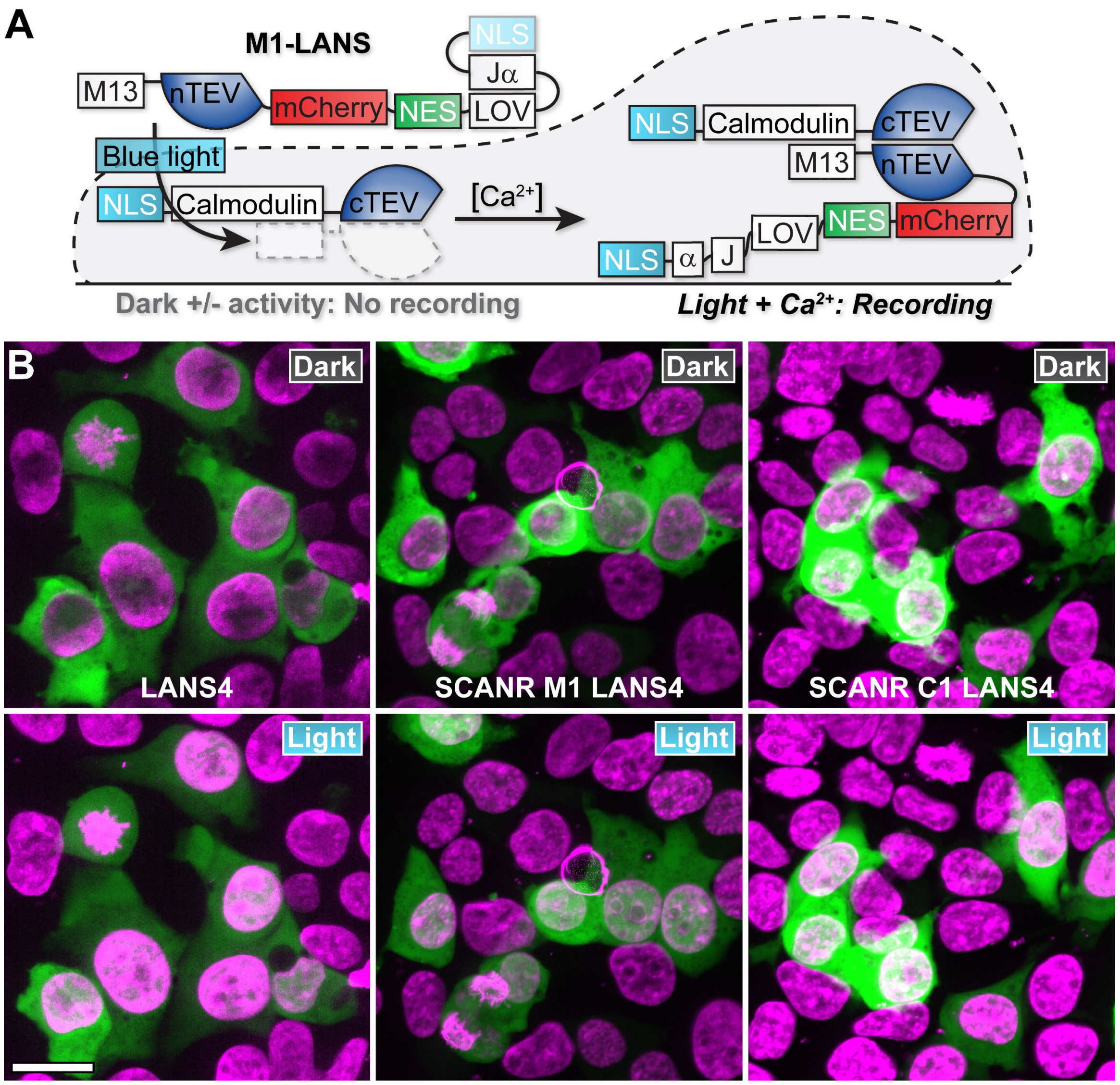

**Figure S2: Application of LANS-based control over subcellular localization to the SCANR system.** (**A**) Schematic depiction of the LANS-modified SCANR constructs. In the dark, the LANS-modified SCANR construct is designed to be sequestered in the cytoplasm, similar to the LiNUS construct. Only in the presence of both blue light and high $Ca^{2+}$ can both partners meet to reform an active TEV protease. (**B**) Expression and illumination of LANS modified constructs. HEK293T cells were transfected with the indicated construct and illuminated with 488 nm light to induce cytoplasm-to-nucleus translocation. The control plasmid, LANS4 shows clear movement of the mCherry fluorescent protein upon the addition of blue light. Neither the M1 nor the C1 -LANS modified constructs showed significant movement into the nucleus after illumination. All images are maximum intensity z-projections of confocal microscopy images. Scale bar = 20 μm.

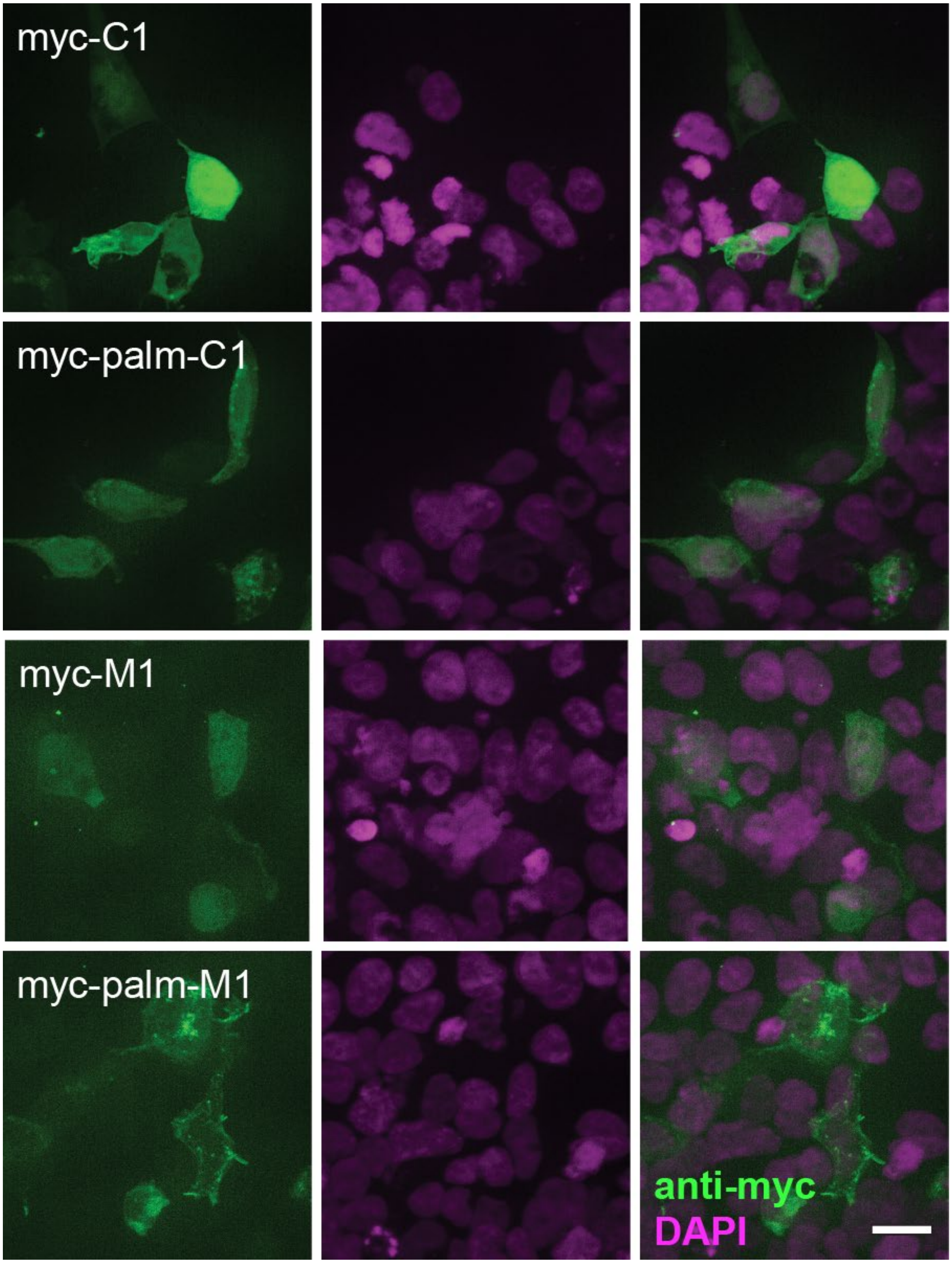

**Figure S3: Expression comparison between palmitoylated (palm) and undirected constructs.** HEK293T cells were transiently transfected with either the palmitoylated or undirected constructs. Using the incorporated myc-tag it was clear that both C1 and M1 express homogeneously throughout the cell. For the palmitoylated constructs, there is a greater anti-myc labelling at the cell membrane. This confirms the correct subcellular localization of these constructs. All images are maximum intensity z-projections of confocal microscopy images. Scale bar = 20 µm

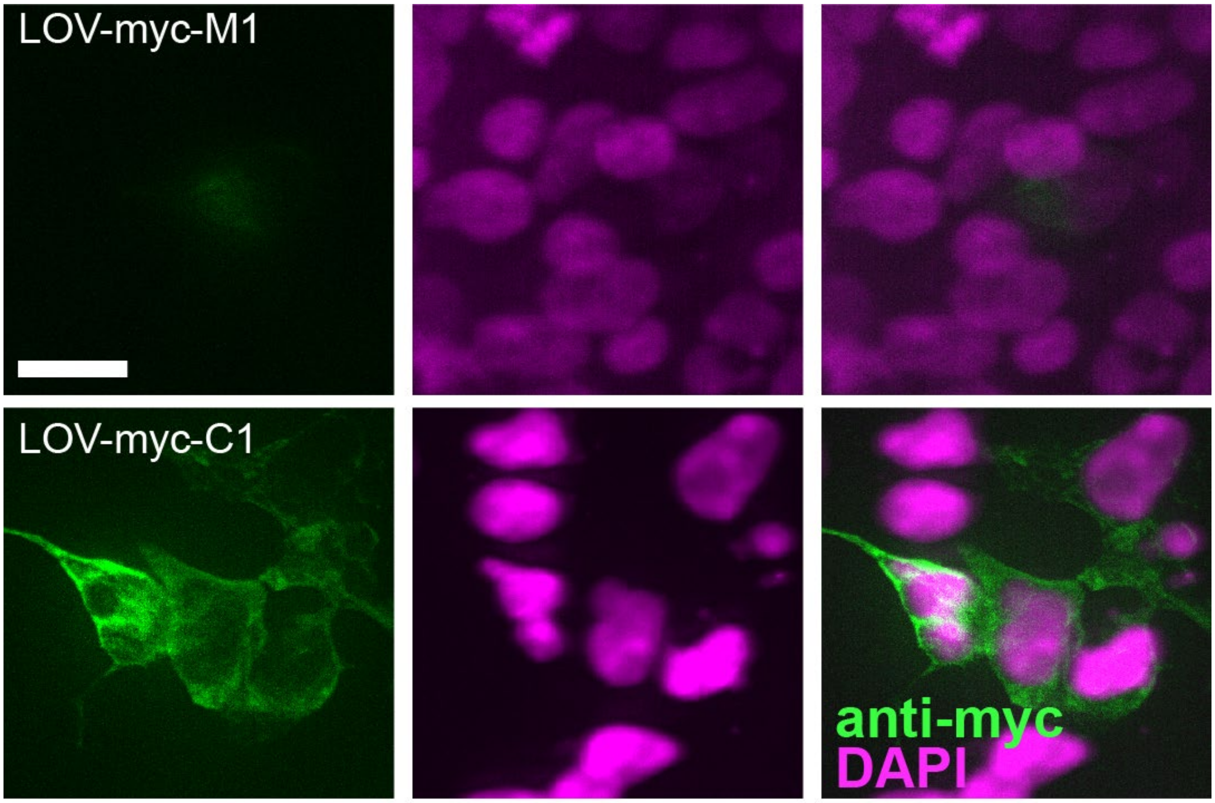

**Figure S4**: **Expression of LOV constructs.** HEK293T cells were transiently transfected with either LOV myc M1 or LOV myc C1and probed with an antibody directed against their appended myc tag. All images are maximum intensity z-projections of confocal microscopy images. Scale bar = 20 μm

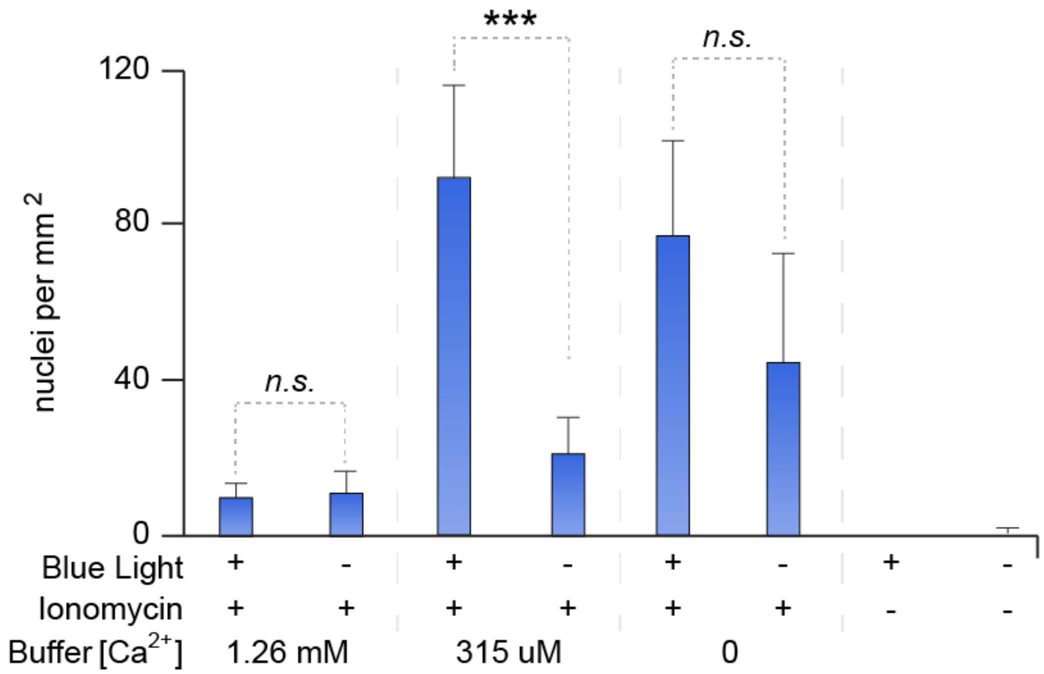

**Figure S5. Characterization of LOV system with changing external buffer $Ca^{2+}$ concentration.** The LOV modified SCANR was transiently transfected into HEK293T cells with the dsRed substrate to evaluate the effectiveness of the system under different conditions. The negative conditions with only blue light or without blue light or ionomycin showed little to no signal. Interestingly, we saw that only an external $Ca^{2+}$ concentration of 315 uM was there a significant difference in the ionomycin only control and ionomycin treated in the presence of blue light. Error bars represent standard deviation. *** $p < 0.001$

**Table S1**: **Primers used for SCANR LEXY modification cloning.** All primers written in the 5' to 3' direction.

| Target Construct | | Amplicon | DNA Oligonucleotide Sequence | |
|---|---|---|---|---|
| C1 LEXY | 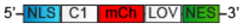 | C1 | *For* | GTGAAGCTGGACGACCAACTGACTGAAGAGCAGATCGCAG |
| | | | *Rev* | CCGCTGCCGCCCATGAAAACTTTATGGCCCCCCCACAATAC |
| | | Linker | *For* | CCATAAAGTTTTCATGGGCGGCAGCGGCGGCAGCGGCGGC |
| | | | *Rev* | CCTCCTCGCCCTTGCTCACGCCGCCGCTGCCGCCGCTGCCGCC |
| | | lin mCh LEXY* | *For* | TCAGTCAGTTGGTCGTCCAGCTTCACGCGCTTTGCTGCGGC |
| | | | *Rev* | GGCAGCGGCGGCGTGAGCAAGGGCGAGGAGGATAACATGGC |
| M1 LEXY | 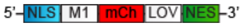 | M1 | *For* | GCGTGAAGCTGGACGGAGAAAGTTTGTTTAAGGGGCCGCGTG |
| | | | *Rev* | GCCGCTGCCGCCCGAGCTTGACAGCCGACCTATAGCTCTG |
| | | Linker | *For* | GCTGTCAAGCTCGGGCGGCAGCGGCGGCAGCGGCGGC |
| | | | *Rev* | CCTCCTCGCCCTTGCTCACGCCGCCGCTGCCGCCGCTGCCGCC |
| | | lin mCh LEXY | *For* | CAAACTTTCTCCGTCCAGCTTCACGCGCTTTGCTGCGGCC |
| | | | *Rev* | GGCAGCGGCGGCGTGAGCAAGGGCGAGGAGGATAACATGGC |
| C1 EGFP LEXY | 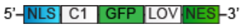 | C1 | *For* | GTGAAGCTGGACGACCAACTGACTGAAGAGCAGATCGCAG |
| | | | *Rev* | CCGCTGCCGCCCATGAAAACTTTATGGCCCCCCCACAATAC |
| | | Linker | *For* | CCATAAAGTTTTCATGGGCGGCAGCGGCGGCAGCGGCGGC |
| | | | *Rev* | CTCCTCGCCCTTGCTCACGCCGCCGCTGCCGCCGCTGCCGCC |
| | | EGFP | *For* | GGCAGCGGCGGCGTGAGCAAGGGCGAGGAGCTGTTCACCG |
| | | | *Rev* | CTCCAGATCCACCCTTGTACAGCTCGTCCATGCCGAGAGTG |
| | | lin mCh LEXY | *For* | TCAGTCAGTTGGTCGTCCAGCTTCACGCGCTTTGCTGCGGC |
| | | | *Rev* | GCTGTACAAGGGTGGATCTGGAGGTTCAGGTGGAAGTTTG |
| C1 LEXY GGS | | | *For* | ACTACCGCCGTCCAGCTTCACGCGCTTTGCTGCGGCCATGC |
| | | | *Rev* | GGCGGTAGTGACCAACTGACTGAAGAGCAGATCGCAG |

**Table S2**: **Primers used for SCANR LINUS and LANS modification cloning.** All primers written in the 5' to 3' direction.

| Target Construct | | Amplicon | | DNA Oligonucleotide Sequence |
|---|---|---|---|---|
| M1 LINUS 9-11 | 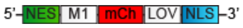 | M1 | *For* | GGTCTTGATATCATGGGAGAAAGTTTGTTTAAGGGGCCG |
| | | | *Rev* | CGCCCTTGCTCACCGAGCTTGACAGCCGACCTATAGCTC |
| Linearized vectors for LINUS cloning | | | *For* | CTTTCTCCCATGATATCAAGACCTGCTAATTTCAAGGC |
| | | | *Rev* | CGGCTGTCAAGCTCGGTGAGCAAGGGCGAGGAGGATAACAT GG |
| M1 LANS1 | 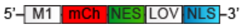 | M1 | *For* | GAGATATACCATGGGAGAAAGTTTGTTTAAGGGGCCGCGTG |
| | | | *Rev* | CTTGCTCACCATCGAGCTTGACAGCCGACCTATAGCTCTGA C |
| Linearized vector for LANS cloning | | | *For* | CTTTCTCCCATGGTATATCTCCTTTGATTGTAAATAAAATG |
| | | | *Rev* | CTGTCAAGCTCGATGGTGAGCAAGGGCGAGGAGGATAAC |

**Table S3**: **Primers used for SCANR CRY2-C1B1 modification cloning.** All primers written in the 5' to 3' direction.

| Target Construct | | Amplicon | DNA Oligonucleotide Sequence | |
|---|---|---|---|---|
| CRY2 dsRed sub | 5'–CRY2–dsR–3' | dsRed sub | *For* | GAACCCTCCCGGGTCGACACCATGGAATTCACCATGGCCGG |
| | | | *Rev* | GGTACGCGGCCGCCTACTTATCGTCGTCATCCTTGTAATC |
| CRY2 mCh C1 | 5'–CRY2–mCh–C1–3' | C1 | *For* | GCAGCAGCCCGGGAGATGGAGCAGAAGCTGATCTCAGAGG |
| | | | *Rev* | CTCGAGCGGCCGCGGTACCTCACATGAAAAC |
| CRY2 mCh M1 | 5'–CRY2–mCh–M1–3' | M1 | *For* | GCAGCAGCCCGGGAGATGGAGCAGAAGCTGATCTCAGAGG |
| | | | *Rev* | CTCGAGCGGCCGCGGTACCTCACGAGC |
| palm C1 | 5'–p–C1–3' | C1 | *For* | GATGAGGACCAAAAGATCATGGAGCAGAAGCTGATCTCAGAGG |
| | | | *Rev* | GGGAGAGGGGCGGAATTCTTACATGAAAACTTTATGGCCCCCCC |
| | | lin C1B1* | *For* | GATCTTTTGGTCCTCATCATTCTTTTCAAC |
| | | | *Rev* | CGAGCGGCCGCGGTACCTCACCATGAAAACTTTATGGCC |
| palm M1 | 5'–p–M1–3' | M1 | *For* | GATGAGGACCAAAAGATCATGGAGCAGAAGCTGATCTCAGAGG |
| | | | *Rev* | GGGAGAGGGGCGGAATTCTTACGAGCTTGACAGCCGACCTATAG |
| | | lin C1B1 | *For* | GATCTTTTGGTCCTCATCATTCTTTTCAAC |
| | | | *Rev* | CTCGAGCGGCCGCGGTACCTCAAGTTTGGAAGTTGGTTGTC |

**Table S4**: **Primers used for SCANR 3xFKBP-FRB modification cloning.** All primers written in the 5’ to 3’ direction. Asterisk denotes constructs that required repair by site directed mutagenesis with the indicated primers.

| Target Construct | | Amplicon | | DNA Oligonucleotide Sequence |
|---|---|---|---|---|
| M1 H2B $FKBP_{3x}$ | 5’ – M1 – H2B – 3xFKBP – 3’ | H2B | *For* | TCTAGAATGCCTGAACCCTCTAAGTCTGCTCCAG |
| | | | *Rev* | TCTAGAGGTGGCGACCGGTGGATCCTTAGAGCTAG |
| M1 H2B $FKBP_{3x}$ * site-directed mutagenesis to fix frame | | | *For* | GTCAGAGCTATAGGTCGGCTGTCAAGCTCGCTCGAGTCTAG AATG |
| | | | *Rev* | CATTCTAGACTCGAGCGAGCTTGACAGCCGACCTATAGCTC TGAC |
| C1 $FKBP_{3x}$ | 5’ – C1 – 3xFKBP – 3’ | C1 | *For* | GGCTAGCAGAATGGACCAACTGACTGAAGAGCAGATC |
| | | | *Rev* | CAAGCTTCGCTTCCCATGAAAACTTTATGGCCCCCCC |
| | | lin $FKBP_{3x}$ | *For* | CCATTCTGCTAGCCAGCTTGGGTCTCCCTATAG |
| | | | *Rev* | GTTTTCATGGGAAGCGAAGCTTGGTACCGAGCTCGG |
| C1 Gal4 VP16 $FKBP_{3x}$ | 5’ – C1 – G4 – 3xFKBP – 3’ | Gal4 VP16 | *For* | AAGCTTCCGCCCGGGCCATGAAGCTACTGTCTTCTATC |
| | | | *Rev* | TCTAGACCCACCGTACTCGTCAATTCCAAGGGCATCGG |
| C1 3xNLS $FKBP_{3x}$ | 5’ – C1 – NLS – 3xFKBP – 3’ | 3x NLS | *For* | AAGCTTCCAATGATCCAAAAAAGAAGAG |
| | | | *Rev* | CTAGATCCGGTGGATCCTACCTTTCTC |
| 6x His FRB NES | 5’ – 6H – FRB – NES – 3’ | FRB | *For* | CATCATCATGGTATCCTCTGGCATGAGATGTGG |
| | | | *Rev* | CGCGGCCGCCTACTTTGAGATTCGTCGGAACACATG |
| | | lin 6x His NES | *For* | CCAGAGGATACCATGATGATGATGATGATGAGAACC |
| | | | *Rev* | GACGAATCTCAAAGTAGGCGGCCGCGACTCTAGATC |
| ERT2 C1 | 5’ – ERT – myc – C1 – 3’ | ERT2 | *For* | GAGGACCTGGAATTCGCCGGTGACATGAGAGCTGCC |
| | | | *Rev* | GTCAGTTGGTCCATACTACCGCCGACTGTGGCAGGGAAACC CTC |
| | | lin myc C1 | *For* | CATGTCACCGGCGAATTCCAGGTCCTCCTCTGAGATC |
| | | | *Rev* | CTGCCACAGTCGGCGGTAGTATGGACCAACTGACTGAAGAG CAG |
| ERT2 M1 | 5’ – ERT – myc – M1 – 3’ | ERT2 | *For* | GAGGACCTGGAATTCGCCGGTGACATGAGAGCTGCC |
| | | | *Rev* | CTTTCTCCCATACTACCGCCGACTGTGGCAGGGAAACCCTC TGC |
| | | lin myc M1 | *For* | CATGTCACCGGCGAATTCCAGGTCCTCCTCTGAGATC |
| | | | *Rev* | CCACAGTCGGCGGTAGTATGGGAGAAAGTTTG |

**Table S5**: **Primers used for SCANR LOV cloning.** All primers written in the 5' to 3' direction. Asterisk denotes constructs that required repair by site directed mutagenesis with the indicated primers.

| Target Construct | | Amplicon | | DNA Oligonucleotide Sequence |
|---|---|---|---|---|
| LOV myc C1 | 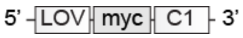 | LOV Jα | *For* | CAAGCTTGCCACCATGGGATCCTTGGCTACTACACTTG |
| | | | *Rev* | CAGCTTCTGCTCAAGTTCTTTTGCCGCCTCATC |
| | | lin myc C1 | *For* | GCCAAGGATCCCATGGTGGCAAGCTTGGGTCTCC |
| | | | *Rev* | GGCAAAAGAACTTGAGCAGAAGCTGATCTCAGAGGAG |
| LOV myc M1 | 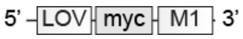 | LOV Jα | *For* | CAAGCTTGCCACCATGGGATCCTTGGCTACTACACTTG |
| | | | *Rev* | CAGCTTCTGCTCAAGTTCTTTTGCCGCCTCATC |
| | | lin myc M1 | *For* | GCCAAGGATCCCATGGTGGCAAGCTTGGGTCTCC |
| | | | *Rev* | GGCAAAAGAACTTGAGCAGAAGCTGATCTCAGAGGAG |
| EGFP LOV myc C1/M1 | 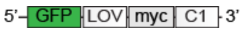 | EGFP | *For* | CCAAGCTTGCCACCATGGTGAGCAAGGGCGAGGAGCTGTTC AC |
| | | | *Rev* | GTAGCCAAGGATCCCATGGACGCCTTGTACAGCTCGTCCAT GCCG |
| | | lin LOV C1/M1 | *For* | GCCCTTGCTCACCATGGTGGCAAGCTTGGGTCTCCCTATAG |
| | | | *Rev* | CTGTACAAGGCGTCCATGGGATCCTTGGCTACTACACTTGA ACG |
| LOV myc C1 p2a EGFP p2a myc M1 | 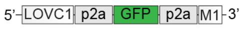 | LOV-Jα | *For* | GGAGGCCCTTCACCATGGGATCCTTGGCTACTACAC |
| | | | *Rev* | GTCAGTTGGTCCATGAATTCCAGGTCCTCCTCTGAGATC |
| | | lin C1 p2a EGFP p2a myc M1 | *For* | GCCAAGGATCCCATGGTGAAGGGCCTCCTCCGGAAATTG |
| | | | *Rev* | GGAGGACCTGGAATTCATGGACCAACTGACTGAAGAGC |

### References

1. Wehr, M. C. *et al.* Monitoring regulated protein-protein interactions using split TEV. *Nat. Methods* **3,** 985–993 (2006).
2. Quan, J. & Tian, J. Circular polymerase extension cloning for high-throughput cloning of complex and combinatorial DNA libraries. *Nat. Protoc.* **6,** 242–51 (2011).
3. O’Neill, B. K. & Laughlin, S. T. Neuronal Calcium Recording with an Engineered TEV Protease. *ACS Chem. Biol.* **13,** 1159–1164 (2018).
